# Dopamine D2R and opioid MOR availability in autism spectrum disorder

**DOI:** 10.1101/2024.04.09.588651

**Authors:** Tuomo Noppari, Jouni Tuisku, Lasse Lukkarinen, Pekka Tani, Nina Lindberg, Emma Saure, Hannu Lauerma, Jari Tiihonen, Jussi Hirvonen, Semi Helin, Johan Rajander, Juha Salmi, Lauri Nummenmaa

**Author notes:** Corrensponding author: Tuomo Noppari, Department of Psychiatry, Helsinki University Hospital, PL 590, 00029 HUS, Helsinki, Finland,. Abnormal opioid-dopamine interaction in autism.

## Abstract

Opioid and dopamine receptor systems are implicated in the pathoetiology of autism, but *in vivo* human brain imaging evidence for their role remains elusive. Here, we investigated regional type 2 dopamine and mu-opioid receptor (D2R and MOR, respectively) availabilities and regional interactions between the two neuromodulatory systems associated with autism spectrum disorder (ASD). In vivo positron emission tomography (PET) with radioligands [11C]raclopride (D2R) and [11C]carfentanil (MOR) was carried out in 16 adult males with high functioning ASD and 19 age and sex matched controls. Regional group differences in D2R and MOR receptor availabilities were tested with linear mixed models and associations between regional receptor availabilities were examined with correlations. There were no group differences in whole-brain voxel-wise analysis of DR2 but ROI analysis presented a lower overall mean D2R availability in striatum of the ASD versus control group. Post hoc regional analysis revealed reduced D2R availability in nucleus accumbens of the ASD group. The whole-brain voxel-wise analysis of MOR revealed precuneal up-regulation in the ASD group, but there was no overall group difference in the ROI analysis for MOR. MOR down-regulation was observed in the hippocampi of the ASD group in a post hoc analysis. Regional correlations between D2R and MOR availabilities were weaker in the ASD group versus control group in the amygdala and nucleus accumbens. These alterations may translate to disrupted modulation of social motivation and reward in ASD.

## Introduction

Autism spectrum disorder (ASD) is a developmental neuropsychiatric disorder with disabling difficulties in social communication and interaction and restricted and repetitive behavior [1]. ASD is strongly heritable with a complex genetic background [2, 3], but its neurochemical basis remains poorly understood. Aberrancies in different neuroreceptor systems [4] have been proposed to underlie ASD, but human studies have yielded only limited evidence for the role of specific neuroreceptor systems. Animal studies and neuropharmacological evidence, however, suggest that the endogenous dopamine and opioid systems might be the key molecular pathways underlying ASD [4–10].

Dopamine is released from the ventral tegmental area (VTA), which projects to the prefrontal cortex (PFC) and nucleus accumbens (NAcc) of the ventral striatum, forming the mesocorticolimbic circuit (MCL) that controls reward and motivated behavior [5, 11]. Dopamine released from substantia nigra pars compacta (SNpc), in turn, projects to the dorsal striatum (DS), forming the nigrostriatal (NS) pathway, controlling goal-directed and habitual behavior [5]. Dopamine receptors are divided into two main groups D1-(D1R, D3R) and D2-receptors (D2R, D4R, D5R), of which D1R and D2R moderate locomotion, reward, reinforcement, learning, and memory. Functional magnetic resonance imaging (fMRI) studies in humans have suggested that the dysfunction of the MCL is associated with ASD [12–15]. Human positron emission tomography (PET) studies have further shown deviant dopamine signaling in ASD compared to controls, but the results have been controversial [16, 17]. There is also support for dopamine dysregulation in ASD from genetic studies [18]. Finally, dopamine antagonist medications such as risperidone and aripiprazole may alleviate autism symptoms, including irritability and stereotypic behavior [4–6, 19].

The endogenous opioid system controls a wide variety of human functions, from basic homeostasis and antinociception to emotions, social attachment, and reward [9, 20–23]. Neuropeptide beta-endorphin is released from hypothalamic nuclei and mesolimbic structures, projecting to other hypothalamic and corticolimbic structures, frontal lobes, brainstem, and periaqueductal grey [9, 20]. Beta-endorphin binds to three different opioid receptors, mu-(MOR), delta-(DOR), and kappa-(KOR), which are differentially distributed across the cortical and subcortical areas. Since the late seventies, opioid system dysfunctions have been proposed to be linked with autism [7, 9, 24, 25], and more recently, MOR knock-out mice were shown to manifest autism-related behaviors [10]. In animals, μ-opioid agonists also influence the social approach and withdrawal behavior depending on the dose, duration of dosing, and the social context [8]. Clinical studies in humans have shown that opiate addicts under low-dose opiate maintenance treatment display deficits of social interaction and social cognition with autism-like behaviors [8, 26–28]. Moreover, opioid antagonists may improve hyperactivity and restlessness symptoms in autism [4, 29]. These endogenous opioid system aberrancies in ASD have not been validated with any human imaging studies.

The interaction of the dopamine and opioid systems is critical for reward processing [30– 33]. In animal microdialysis studies, mu-opioid agonists in the VTA increased the basal level of dopamine in the striatum, whereas antagonists decreased it [34, 35]. In human [11C]carfentanil PET studies, the dopaminergic drug amphetamine has raised mu-opioid levels in the striatum, frontal cortex, thalamus, insula, and anterior cingulum (ACC) and cerebellum [36, 37]. Multiligand PET studies have confirmed that D2R and MOR availabilities correlate positively with each other in the striatum, [38, 39], and that this linkage may be disrupted in specific conditions such as morbid obesity [39].

Here, we used in vivo PET with radioligands [11C]raclopride (binding to D2Rs) and [11C]carfentanil (binding to MORs) to compare regional receptor availabilities and the interactions between the receptor systems in ASD individuals and controls. We found that ASD is associated with striatal D2R basal availability downregulation and disruption of D2R and MOR interaction.

## Materials and methods

### Participants

Sixteen individuals with high-functioning ASD (mean age 30 years, range 23–42 years) and 19 controls (mean age 29 years, range 21–49 years) participated in the study. All participants gave informed, written consent and were compensated for their participation. The ethics board of the Hospital District of Southwest Finland approved the protocol, and the study was conducted in accordance with the Declaration of Helsinki. The study also obtained a research permit from the Hospital District of Helsinki and Uusimaa since the ASD group participants were largely recruited from this area. General inclusion criteria for both groups were male sex and age of 20–50 years. General exclusion criteria for both groups were current use of narcotics, current abusive use of alcohol (over 14 units per week), severe axis I psychiatric illnesses, autoimmune illnesses, current medical conditions, and the standard MRI contraindications. Structural brain abnormalities that are clinically relevant or could bias the analyses were excluded by a consultant neuroradiologist.

An additional inclusion criterion for the ASD group was a valid autism spectrum disorder diagnosis (DSM-5 299.00). Participants in the ASD group were volunteers from the Helsinki and Turku University Hospital Neuropsychiatric Clinics, one participant was also recruited from Neuropsychiatric Clinic Proneuron, in Espoo. Diagnoses were verified by a neuropsychologist (JS), neurologist (TN), and psychiatrist (PT) following DSM-5 criteria based on patient history, all accessible information from birth records, well-baby clinics, and school healthcare. Autism Diagnostic Observation Schedule-Second Edition (ADOS-2) assessment [40] was also performed by trained, research-reliable clinical psychologist (ES) to clarify the autism diagnostics. Possible opioid and dopaminergic medications were paused during measurements. Clinical information and characteristics of the ASD participants are found in **Supplement S1**.

Additional inclusion criteria for the control group were no history of neurological or psychiatric disorders. The control participants were screened for medical conditions from their patient histories and their use of prescribed medication was double-checked from the Finnish medical database. Clinical information and characteristics of the control participants are found in **Supplement S1**.

All participants completed the Autism Spectrum Quotient (AQ) questionnaire for the evaluation of ASD -related traits [41, 42]. None of the participants had previous or current severe mental disorders based on the SCID-I interview [43]. The education of all the participants was classified into three different categories: (1) primary school, (2) secondary degree, (3) university degree. All the participants were also screened for their clinical status, with comprehensive general blood laboratory exams, including a urinary test for the use of narcotics.

### Data acquisition

PET imaging was carried out with General Electric (GE) Healthcare Discovery VCT and Discovery 690 PET/CT scanners in Turku PET center. MOR availability was measured with high-affinity agonist radioligand [11C]carfentanil [44] and D2R availability with high-affinity antagonist radioligand [11C]raclopride [45]. Radiotracer synthesis has been described previously [46, 47] Both radioligands were administered as a rapid bolus injection, after which the radioactivity in the brain was measured for 51 minutes. Injected doses were 245 ± 11 MBq for [11C]carfentanil and 254 ± 11 MBq for [11C]raclopride). The [11C]carfentanil and [11C]raclopride PET imaging were performed on the same day > 2.5 hours apart. All PET images were reconstructed using 13 timeframes (3 x 1min, 4 × 3 min, 6 × 6 min; total of 51 minutes). High-resolution anatomical T1-weighted images (TR 9.8 ms, TE 4.6 ms, flip angle 7º, 250 mm FOV, 256 × 256 voxel matrix with 1 mm^3^ isotropic voxel size) were obtained with 3T PET-MRI scanner (Philips Ingenuity TF PET-MR device, Philips Healthcare) for reference and normalization purposes.

### PET preprocessing and modelling

PET images were preprocessed in MATLAB (The MathWorks, Inc., Natick, Massachusetts, United States) using Magia pipeline [48] (https://github.com/tkkarjal/magia), which utilizes SPM12 (The Wellcome Trust Centre for Neuroimaging, Institute of Neurology, University College London, https://www.fil.ion.ucl.ac.uk/spm/software/spm12) in the PET data motion correction and image registration and FreeSurfer (version 6.3, https://freesurfer.net) in an automated region of interest (ROI) delineation. Six bilateral ROIs including the amygdala, thalamus, caudate, globus pallidus (GP), NAcc, and putamen were extracted for D2R from MRI by using FreeSurfer. Additionally, for MOR, we determined seven other ROIs (dorsal anterior cingulate cortices (dACC) and rostral anterior cingulate cortices (rACC), OFC, posterior cingulate cortex (PCC), insula and hippocampus) involved in socioemotional processing [49, 50] and with high MOR binding. Reference tissues volumes (cerebellum for [11C]raclopride and occipital cortex for [11C]carfentanil) were automatically adjusted to account for specific radioligand binding, as described previously [48].

Specific regional binding of [11C]raclopride and [11C]carfentanil was quantified as binding potential relative to non-displaceable binding (*BP*ND) using the simplified reference tissue model (SRTM) [51]. Parametric *BP*ND images were also calculated for voxel-level analysis with basis function implementation of SRTM (bfSRTM) with 300 basis functions. Lower and upper bounds for theta parameter were set to 0.082 1/min and 0.6 1/min for [11C]raclopride and 0.06 1/min and 0.6 1/min for [^11^C]carfentanil. Before the parametric image calculation, the dynamic PET images were smoothed using a Gaussian kernel with 4 mm full width at half maximum to reduce the effect of noise in voxel-level bfSRTM fit. The resulting parametric images were further normalized into MNI152 space and smoothed again using a Gaussian 4 mm filter.

### Statistical analysis

Full-volume analysis of the normalized and smoothed [11C]raclopride and [11C]carfentanil images was carried out with SPM12 using a two-sample Student’s t-test, while controlling for scanner type. [11C]carfentanil analysis included the whole brain volume, whereas voxel-wise [11C]raclopride analysis was restricted to striatum, as regional BP_ND_ for this ligand can be reliably estimated only within these subregions [52]. The results were corrected for multiple comparisons by using false discovery rate (FDR) at p < 0.05.

Statistical analyses for the ROI data were performed with R statistical software version 3.5.1 (The R foundation, Vienna, Austria) using linear mixed-effects models (LMMs). We first tested for overall significant differences between the groups by including group as a fixed factor and participant, ROI, and scanner type as random effects, allowing their intercepts to vary freely. Before the LMM analysis, all *BP*ND values were log-transformed to ensure normality. Normality of the data was evaluated with the Shapiro-Wilk test and checked visually with density and Q-Q plots. We then set up post-hoc analysis to assess group differences in each ROI separately by specifying group as a fixed factor and scanner type as a random effect in LMM. The association between AQ scores and regional radioligand availability was analyzed with LMM, with AQ as a fixed factor and scanner type as a random effect.

Associations between regional [11C]carfentanil and [11C]raclopride *BP*ND were estimated using Pearson correlations. Group differences in the head-to-head regional correlations were tested using Fisher’s R-Z test, and Mantel’s test was used to examine the similarity between correlation structures of MOR / D2R across regions in both groups (i.e., correlation matrices)

### Meta-analysis

To summarize the current data on striatal dopamine PET imaging studies in ASD we conducted a meta-analysis. A PubMed search was conducted on Oct. 3^rd^, 2023, using the keywords “autism spectrum disorder”, “PET”, “positron emission tomography” and “dopamine”. Moreover, a complementary search with Google Scholar was conducted to confirm that no studies were missing. Finally, we screened three of the latest review articles on the issue to ensure the full coverage of our research [16, 53, 54]. Altogether, we found nine potentially eligible articles [55–63], but only six included requisite data on striatal ROIs. Two of these six articles used the same data, so only one was included. Together with our results presented here a total number of six articles were included in the meta-analysis. All the articles were evaluated by three independent researchers of our group (TN, LN, JT) for inclusion in the meta-analysis. There was only one previous study using [11C]raclopride tracer [62] measuring D2R availability, whereas the other studies used [18F]FDOPA [56, 58, 60, 61] measuring presynaptic dopamine synthesis capacity or D1R binding [11C]SCH23390 [57] as tracers (see **Supplement S2** for more details). As one of the articles presented required data only in their supplemental figure and it was also not readily accessible from the authors, we used plotdigitizer tool to extract baseline values from the accessible figure [62]. Based on reported striatal group means, standard deviations and sample sizes we calculated the standardized mean difference (SMD, Cohen’s d) for each study and a pooled sample size weighted SMD with R software metafor package.

## Results

Mean group-wise [^11^C]raclopride and [^11^C] carfentanil binding maps are shown in **Figure 1**. Group-wise regional binding potentials for the radioligands are shown in **Figure 2**.

**Figure 1.**
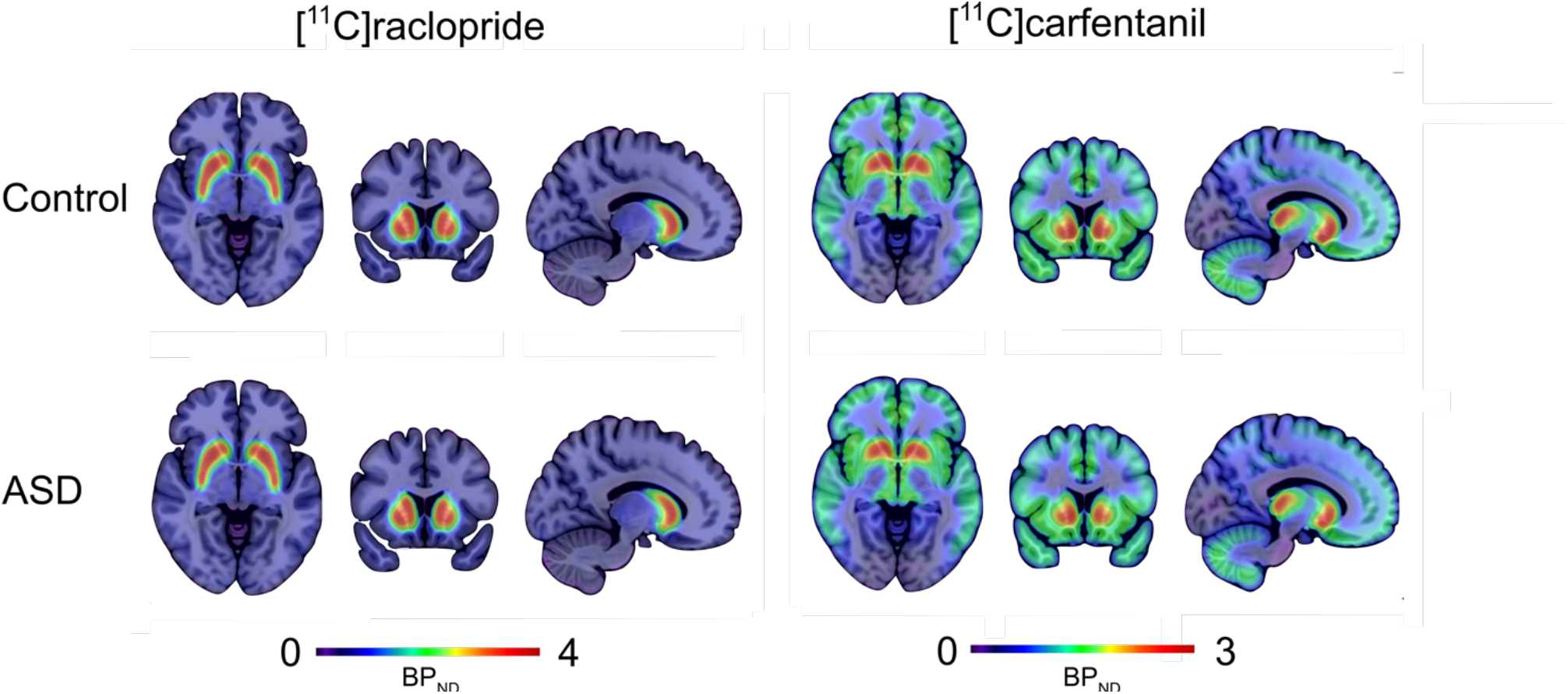
Mean [11C]raclopride and [11C]carfentanil *BP*_ND_ in the control and ASD groups.

**Figure 2.**
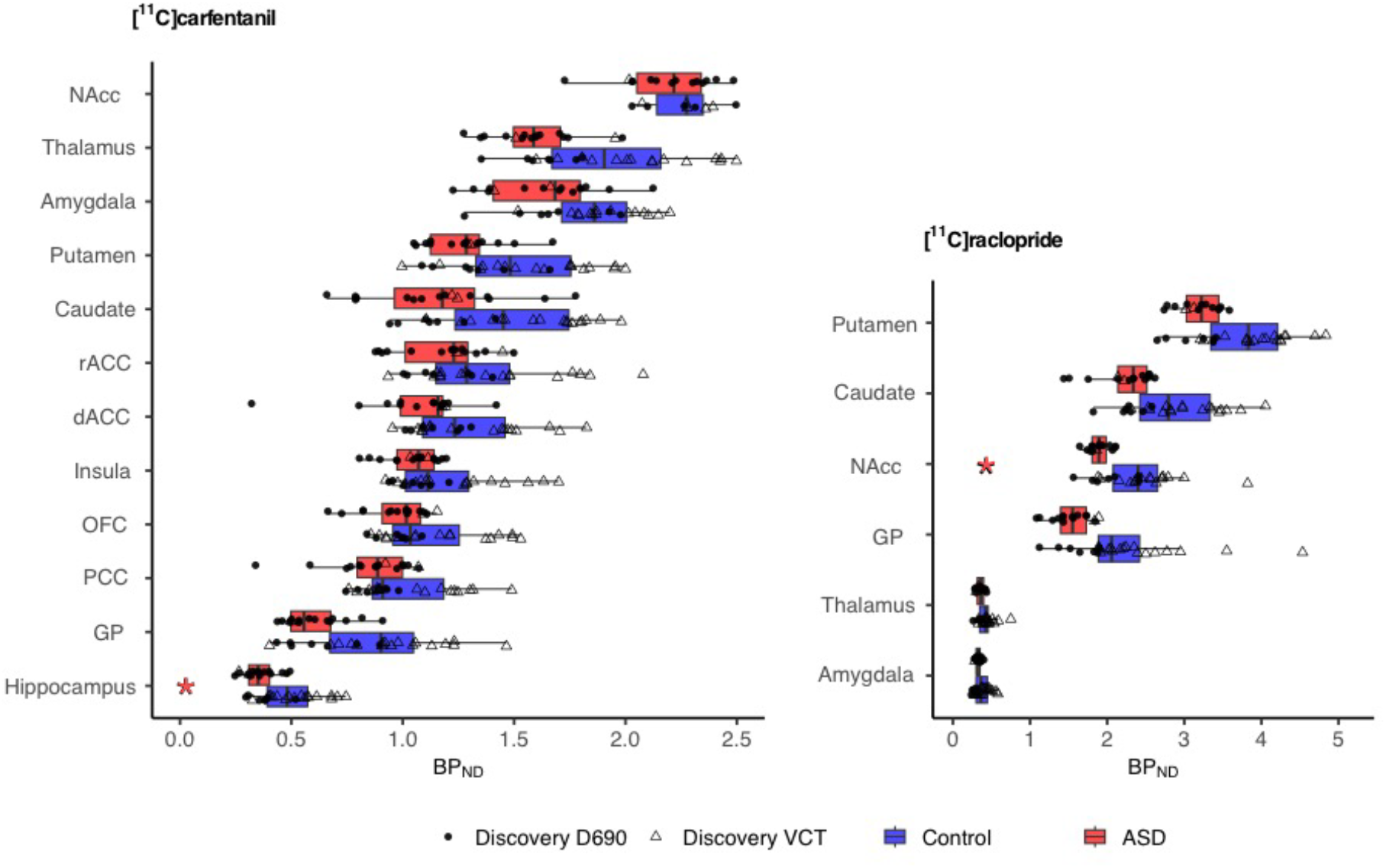
Group-wise binding potentials of [11C]raclopride and [11C]carfentanil *BP*_ND_ in the predefined ROIs. The blue and red boxes mark the 2^nd^ and 3^rd^ quartiles, with the crossline marking the median of the data. Upper and lower whiskers mark 1.5 interquartile ranges from the box. Data with the two different scanners is marked with circles (Discovery 690) and triangles (Discovery VCT). Significant differences between the groups are marked with an asterisk.

**Figure 3.**
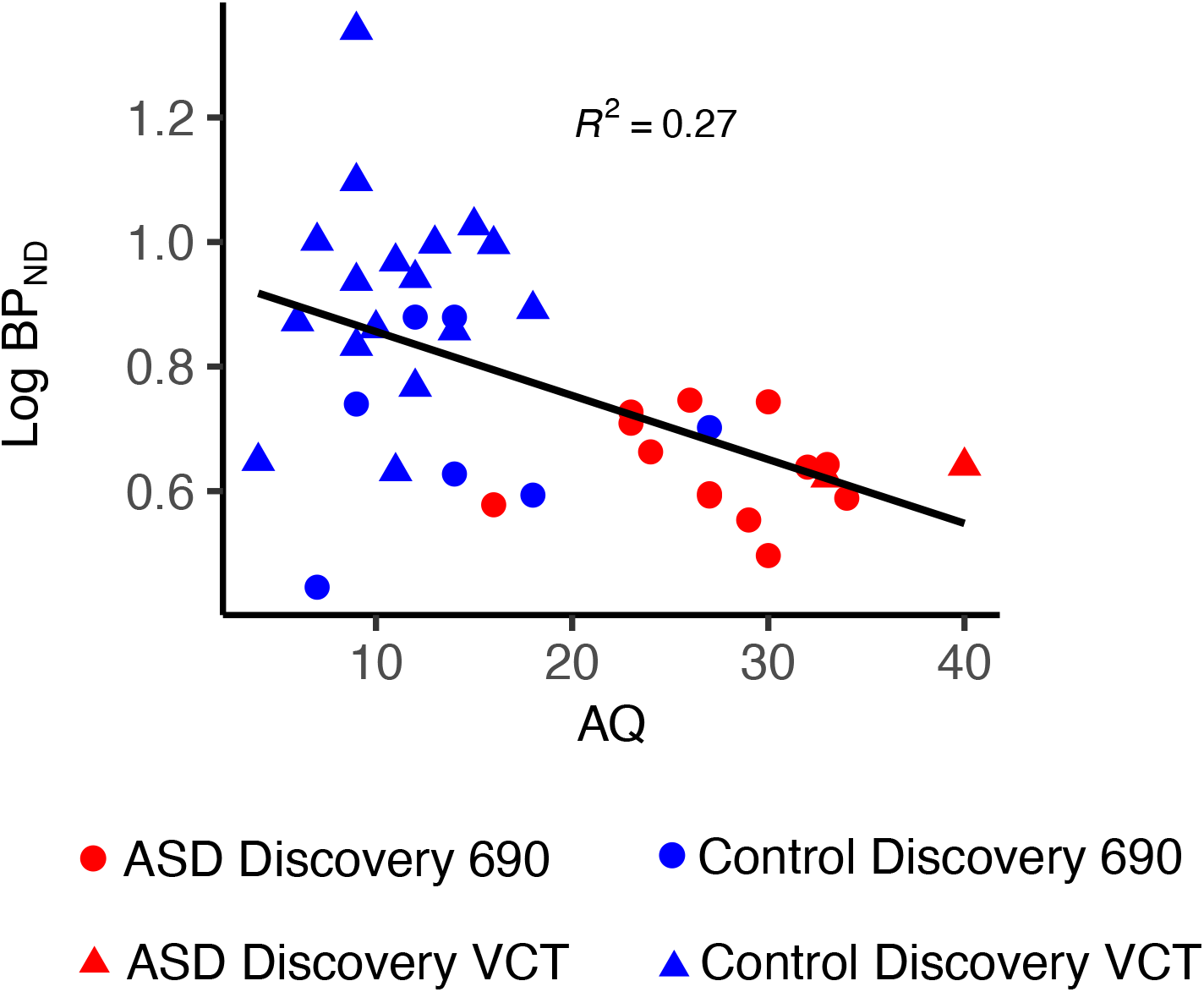
Association and regression line between AQ scores and [11C]raclopride *BP*_ND_ in the NAcc across the ASD and control groups. Data with the two different scanners is marked with circles (Discovery 690) and triangles (Discovery VCT).

**Figure 4.**
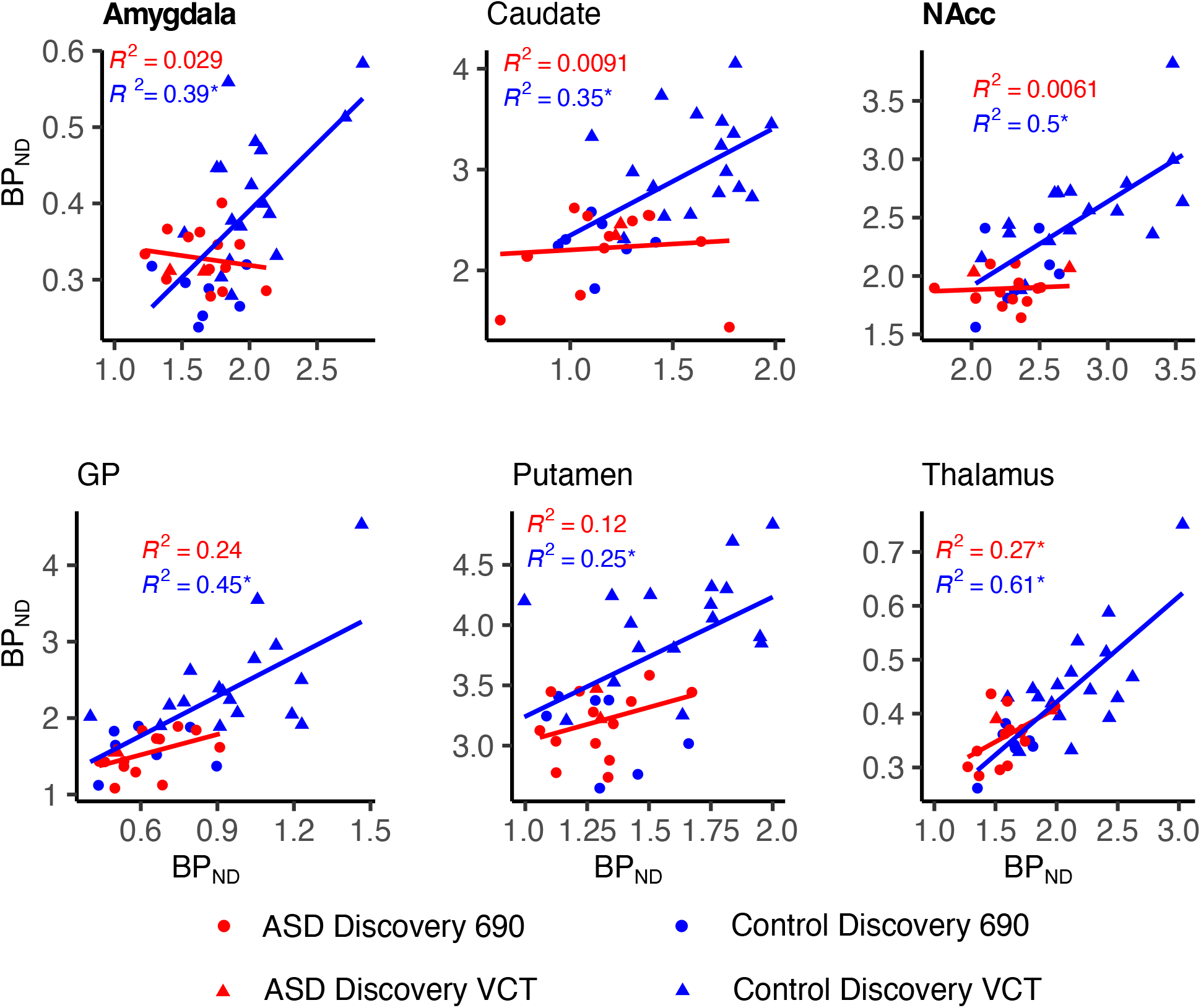
Regional association and LS-regression lines between [11C]carfentanil and [11C]raclopride *BP*_ND_ in control and ASD groups. Significant within-group correlations are marked with an asterisk, and significant differences in correlations between groups (Amygdala and NAcc) with boldface.

**Figure 5.**
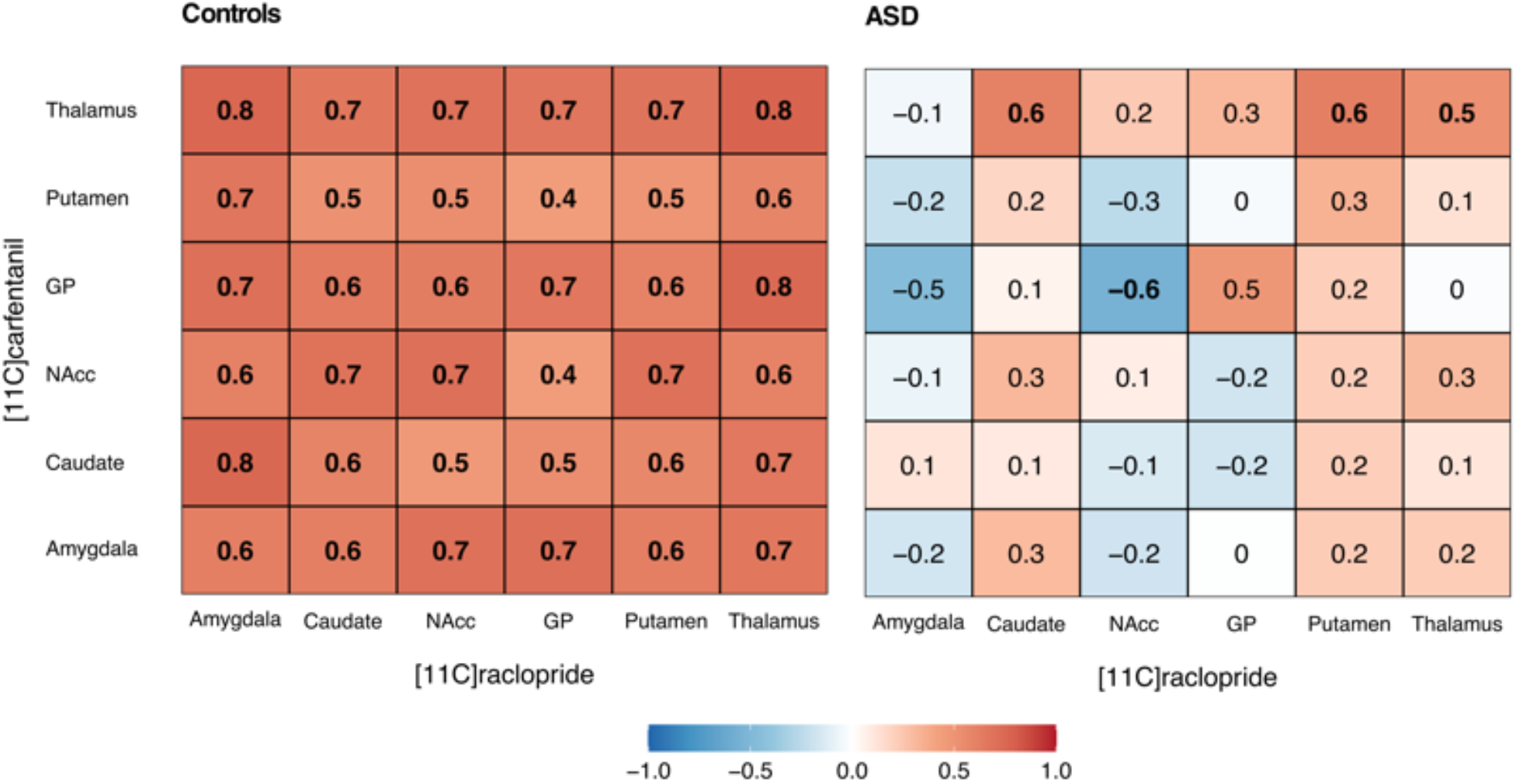
Matrices showing interregional Pearson correlations between [11C]carfentanil (rows) and [11C]raclopride (columns) *BP*_ND_ in the control and ASD groups. Significant correlations are marked with boldface.

### D2R system

Whole-brain voxel-wise analysis did not reveal any significant group differences with [11C]raclopride. The analysis with striatal ROIs revealed significant overall downregulation in D2R availability of the ASD group (p = 0.04). A complementary analysis, including extra-striatal regions thalamus and amygdala, revealed no group differences. Separate regional post-hoc analyses showed that the effect was significant in the NAcc (Estimate 0.135, p = 0.02, Shapiro p = 0.11). *BP*ND in the NAcc was also negatively correlated with AQ scores across all participants (Estimate -0.006, p = 0.033, Shapiro p = 0.181), but this effect was not significant when the groups were analyzed separately.

### MOR system

Whole brain voxel-wise analysis revealed higher *BP*ND in ASD participants compared to controls in the precuneus (p < 0.05, FDR corrected). ROI analysis revealed no overall differences between groups in MOR *BP*ND. However, regional post-hoc analyses indicated MOR downregulation in the ASD group in the hippocampus (Estimate 0.215, p = 0.022, Shapiro p = 0.381). Furthermore, MOR *BP*ND did not correlate with AQ results in any of the ROIs across all participants or within either group separately.

### Dopamine-opioid interaction

In the control group, the *BP*ND for [11C]carfentanil and [11C]raclopride correlated positively across all six ROIs that were analyzed with both ligands. In the ASD group, significant correlation was only observed in thalamus. The correlations were significantly stronger in the control versus ASD group in amygdala (z = 2.48 two-tailed p = 0.013) and NAcc (z = 2.230, two-tailed p = 0.026). The Mantel test did not indicate similarity between the MOR / D2R correlation matrices for control and ASD group (r = 0.65, p = 0.16).

### Meta-analysis of PET-imaging data on dopamine systems in ASD

A fixed effect model showed a standardized mean difference (SMD) of -0.32 (-0.62, -0,03) with heterogeneity of 81.2% (p < 0.001), and a random model showed SMD of -0.25 (-1.06, 0.57) with heterogeneity of 85,6 % (p < 0.001). We found a small overall effect of decreased dopamine tracer binding in ASD compared to controls (Figure 6).

**Figure 6.**
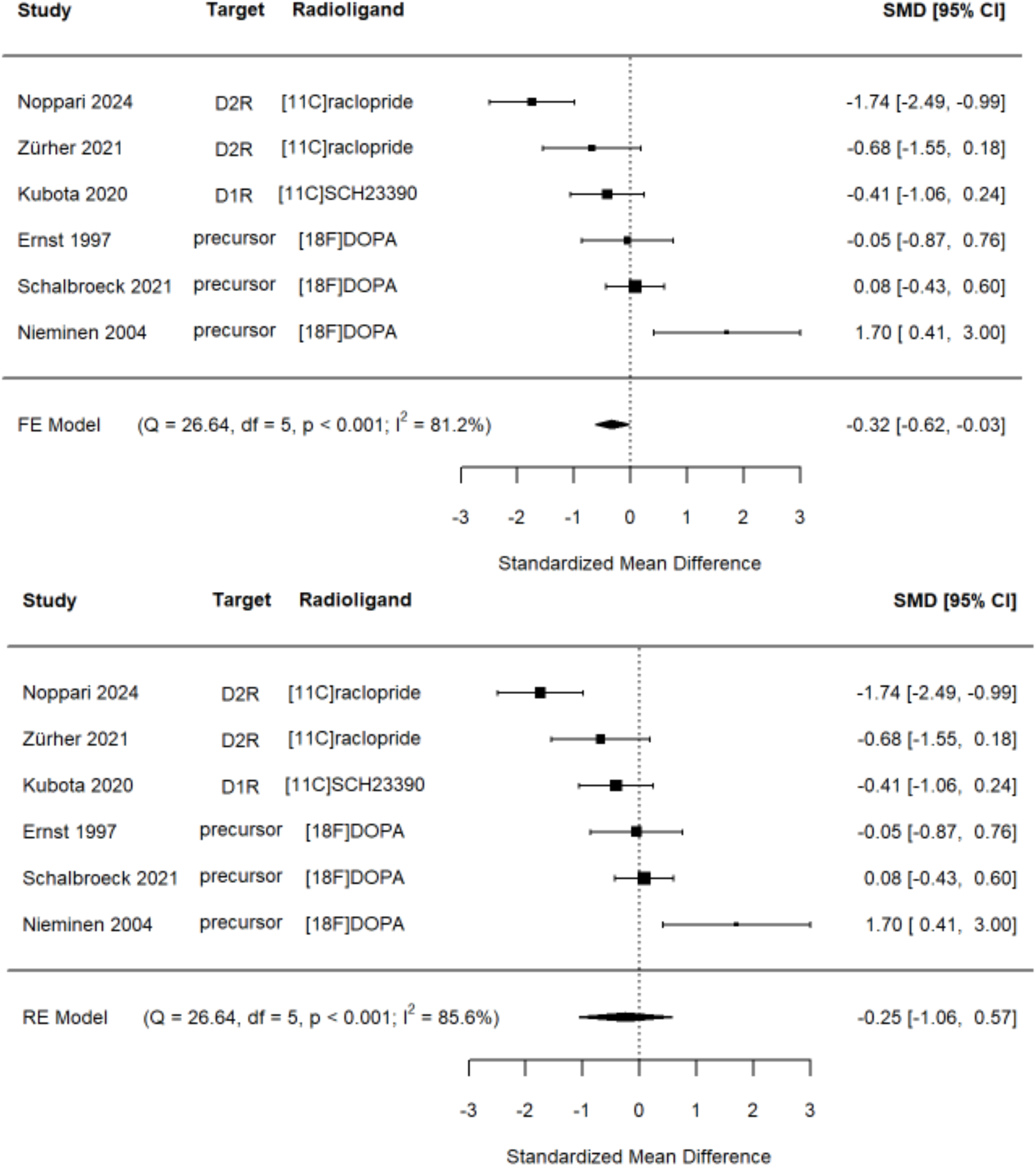
Separate and pooled standardized mean differences (SMDs) in a meta-analysis of six striatal dopamine PET imaging studies with fixed (FE) and random effect (RE) models. Q = Cochran’s Q statistic, df = degrees of freedom, I^2^ = inconsistency index.

## Discussion

Our main findings were that ASD is associated with downregulated striatal D2R availability and aberrant interaction between D2R and MOR systems. Alterations in the MOR system were restricted to the hippocampus and precuneus. Altogether these data suggest that the dopaminergic system might be the main neurotransmitter pathway linked with autism, also partly due to its aberrant interaction with the endogenous MOR system.

### D2R system

Mean *BP*ND for [11C]raclopride in the striatum was lower in the ASD compared to the control group. Our meta-analysis of the dopamine PET studies showed a consistent but a small and heterogeneous overall effect of dopamine downregulation in ASD compared to controls. The studies using FDOPA reported either no differences or dopamine upregulation in the ASD compared to controls, while the other studies using [11C]raclopride and [11C]SCH23390 reported findings similar to ours. Differences in receptor availability, rather than dopamine synthesis capacity (measured by FDOPA), may thus be altered in ASD [64]. Striatal dopamine modulates social and nonsocial reward and motivation [5, 11], which may be disrupted in ASD, according to fMRI findings [13]. In animal studies, the striatal dopaminergic effects have been associated with stereotypic repetitive behavior [65], but the human imaging findings in ASD have not been conclusive [66].

The ROI analysis revealed significantly lower *BP*ND in the NAcc in the ASD group than in the control group, and the AQ scores were also correlated with *BP*ND in the NAcc across all participants. In human fMRI studies, NAcc has been associated with responses to reward and punishment [67], and in animal studies, dopamine level in NAcc has been associated with prosocial behavior (increased release) or social avoidance in defeat situations (decreased release) [68]. Moreover, fMRI studies have shown NAcc hypoactivation during monetary reward anticipation, and in ASD, NAcc is hypoconnected with the posterior cingulate cortex [12, 69, 70]. The present D2R findings could hence relate, for instance, to aberrant social motivation and potentially explain atypical responses to social reward and high social avoidance in ASD [13, 15, 68, 71]. There are also possible links between NAcc function and stereotypic repetitive behavior in ASD [65, 66]. However, none of these findings can be verified within the scope of this study, and further research is needed.

### MOR system

The whole brain voxel-wise analysis showed a significant group difference in the left precuneus with higher *BP*ND in the ASD group. Precuneus is one of the main areas of the Default Mode Network (DMN) critically involved in social cognition and showing dysfunction in ASD [72–76]. PET studies have shown significant distribution of opioid receptors in the human DMN, and opioids modulate its activity in a dose-dependent manner as the low opioid doses inhibit and the high doses enhance it [20]. Our previous study with a largely overlapping sample revealed lower grey matter volume in the left precuneus of the ASD group [77]. The present study provides novel evidence that besides structural and functional alterations, also MOR functioning is abnormal in the precuneus area in individuals with ASD. This could play a role in explaining the symptom spectrum of ASD.

The post-hoc ROI analyses showed significantly lower *BP*ND in the hippocampus in ASD participants compared to controls. There are fMRI studies demonstrating the involvement of the hippocampus in impaired perception of social emotions, as well as atypical learning, memory, and sensory hypersensitivity in ASD [78]. General ex vivo autoradiography in monkeys and humans has found a low density of MOR in the hippocampus [20]. The hippocampus is also part of the MOR-modulated DMN and has stronger functional connectivity with other DMN regions in ASD compared to controls [78]. Moreover, MOR modulates the hippocampal neurogenesis, mediating a negative opioid effect, whereas a positive effect can be seen in MOR null knock-out mice [79, 80]. The animal models of ASD present an impaired hippocampal neurogenesis, possibly explaining some of the behavioral phenotypes of the disorder [81]. Our findings on MOR availability could relate to these forementioned behavioral impairments and aberrant modulation of the DMN and the possibly impaired hippocampal neurogenesis in ASD.

### D2R and MOR system interaction

The *BP*ND of [11C]carfentanil and [11C]raclopride showed high interligand correlations across the ROIs in the control group, while in the ASD group, significant interligand correlation was found only in the thalamus. Based on the direct comparison of the correlation strengths, the association between the two ligands was significantly stronger in the amygdala and NAcc of the control group compared to ASD group. In animal studies, both neurotransmitters have been associated with the modulation of motivational and rewarding aspects of social interaction [71, 82]. The NAcc has been identified as the key site for this dopamine and opioid modulation of social interaction, and the role of the amygdala has also been recognized in earlier studies [69, 71, 82, 83]. PET studies suggest a largely similar interaction between endogenous opioid and dopamine signaling in humans than observed in animals [36–38]. Interestingly, the interaction of these two neurotransmitters in NAcc is also implicated in the modulation of pain sensations [84, 85] that are both perceived and expressed abnormally in ASD [86]. While the existing knowledge of the aberrant pain modulation in ASD is still scarce [86], the previous and our present findings suggest a possible direction for further investigations.

## Limitations

Despite our modest sample size, we found significant group differences and correlations to ASD symptoms in D2R and MOR systems. A more detailed examination of the role of these two neuromodulatory systems in different ASD symptom domains would require a larger sample size. Using two different PET scanners was suboptimal, but this was accounted for in the statistical analyses. To reduce the heterogeneity in the ASD sample, we included only male participants, and therefore, our results may not be generalizable to the ASD population as a whole [87]. Since [11C]raclopride ligand binds specifically to D2R and reflects receptor availability reliably only in the striatum, our findings should not be generalized to other mechanisms in the dopamine system (e.g., extra-striatal or D1R dopamine binding). Moreover, our measurement was restricted to basal levels of [11C]raclopride and [11C]carfentanil binding. A comparison between basal versus phasic task-related imaging protocols could help obtain more detailed functional interpretation for aberrant D2R and MOR systems in ASD.

## Conclusions

Taken together, our findings reveal that ASD is associated with downregulated D2R availability and aberrant dopamine-opioid interaction in the striatum, as well as deviant MOR availability in the areas related to DMN. These findings pave the way for new avenues to study the neuromolecular pathways leading to ASD symptoms and to develop related pharmacological treatments.

## Supporting information

Table S1, Clinical and sociodemographic characteristics of the participants

Table S2, Imaging methods and participant characteristics for the meta-analysis of striatal dopamine PET studies in ASD.

## Acknowledgements

This study was supported by the Academy of Finland (grant numbers 294897 and 332225 to L.N); Finnish Governmental Research Funding (VTR) for Turku University Hospital (L.N.), Finnish Governmental Research Funding (VTR) for Helsinki University Hospital (T.N.).

## Conflict of Interest

None

## Notes

### Competing Interest Statement

The authors have declared no competing interest.

