## Supplementary material for "Dopamine D2R and opioid MOR availability in autism spectrum disorder": Table S1, Clinical and sociodemographic characteristics of the participants

| Patient | Age | BMI | AQ Score* | ADOS Score | Diagnosis | Medication | Education* | Handedness |
|---|---|---|---|---|---|---|---|---|
| **ASD** | | | | | | | | |
| 1 | 29 | 30 | 33 | 16 | ASD | None | Second degree | Left |
| 2 | 31 | 21 | 26 | 7 | ASD | None | University | Left |
| 3 | 27 | 21 | 33 | 10 | ASD | None | Second degree | Right |
| 4 | 26 | 34 | 40 | 7 | ASD, ADHD | None | Second degree | Right |
| 5 | 31 | 30 | 30 | 10 | ASD, MAD | Fluoxetine | Second degree | Right |
| 6 | 27 | 29 | 23 | 7 | ASD, MAD | Zolpidem (stopped 1 day before) | Second degree | Right |
| 7 | 25 | 30 | 27 | 16 | ASD | None | Second degree | Right |
| 8 | 35 | 41 | 16 | 12 | ASD, MAD | Melatonin | Primary School | Left |
| 9 | 28 | 22 | 32 | 13 | ASD | None | University | Left |
| 10 | 42 | 19 | 23 | 2 | ASD, MAD | Venlafaxine, Mirtazapine | Second degree | Right |
| 11 | 40 | 18 | 34 | 12 | ASD | None | Second degree | Right |
| 12 | 23 | 24 | 21 | 17 | ASD, MAD | Escitalopram | Second degree | Left |
| 13 | 28 | 27 | 24 | 14 | ASD, MAD | None | Second degree | Right |
| 14 | 31 | 19 | 30 | 14 | ASD, ADHD | None | Primary School | Right |
| 15 | 29 | 22 | 27 | 18 | ASD | None | Second degree | Right |
| 16 | 27 | 21 | 29 | 4 | ASD | None | Second degree | Right |
| *Mean (Std.)* | *30 (5)* | *26 (7)* | *28 (6)* | *11 (5)* | | | | |
| **Controls** | | | | | | | | |
| 1 | 30 | 29 | 9 | _- | None | None | Second degree | Right |
| 2 | 21 | 23 | 10 | - | None | None | Second degree | Right |
| 3 | 22 | 25 | 18 | - | None | None | Second degree | Right |
| 4 | 23 | 26 | 15 | - | None | None | Second degree | Right |
| 5 | 21 | 26 | 16 | - | None | None | Second degree | Right |
| 6 | 34 | 23 | 12 | - | None | None | Second degree | Right |
| 7 | 47 | 24 | 12 | - | None | None | University | Left |
| 8 | 23 | 24 | 9 | - | None | None | Second degree | Right |
| 9 | 31 | 25 | 11 | - | None | None | University | Right |
| 10 | 20 | 22 | 14 | - | None | None | Second degree | Right |
| 11 | 43 | 28 | 4 | - | None | None | University | Right |
| 12 | 25 | 27 | 9 | - | None | None | University | Right |
| 13 | 26 | 22 | 6 | - | None | None | University | Left |
| 14 | 28 | 27 | 13 | - | None | None | University | Right |
| 15 | 32 | 22 | 7 | - | None | None | Second degree | Left |
| 16 | 37 | 27 | 11 | - | None | None | University | Right |
| 17 | 33 | 27 | 9 | - | None | None | University | Right |
| 18 | 24 | 25 | 9 | - | None | None | University | Right |
| 19 | 22 | 27 | 14 | - | None | None | Second degree | Right |
| 20 | 22 | 23 | 12 | - | None | None | Second degree | Right |
| 21 | 26 | 24 | 14 | - | None | None | University | Left |
| 22 | 45 | 27 | 18 | - | None | None | Second degree | Right |
| 23 | 49 | 29 | 7 | - | None | None | Second degree | Right |
| 24 | 21 | 26 | 27 | - | None | None | Second degree | Right |
| *Mean (Std.)* | *29 (9)* | *25 (2)* | *12 (5)* | | | | | |

Note: ASD=Asperger´s syndrome, ADHD=Attention-Deficit/Hyperactivity disorder, MAD=Mood and Anxiety Disorder. Statistically significant group differences ($p<0.05$) are marked with an asterisk.
