## Supplementary material for "Dopamine D2R and opioid MOR availability in autism spectrum disorder": Table S2, Imaging methods and participant characteristics for the meta-analysis of striatal dopamine PET studies in ASD.

| Study | Tracer | Target | ASD n | Controls n | Age Mean (Std.) | Diagnosis |
|---|---|---|---|---|---|---|
| *Ernst et al, 1997* | [18F]FDOPA | Precursor | 8 male<br>6 female | 7 male<br>3 female | ASD: 13(2)<br>Controls: 14(2) | DSM-III-R |
| *Nieminen von Wendt et al, 2004* | [18F]FDOPA | Precursor | 8 male | 5 male | ASD: 29(6)<br>Controls: 3(5) | DSV-IV, ICD-10 |
| *Kubota et al, 2020* | [11C]SCH23390 | D1R | 18 male | 20 male | ASD: 33(8)<br>Controls: 30(6) | DSM-IV-TR |
| *Schalbroeck et al, 2021* | [18F]-FDOPA | Precursor | 28 male<br>16 female | 14 male<br>8 female | ASD: 24(3),<br>Controls: 23(2) | ADOS-2 AQ |
| *Zürher et al, 2021* | [11C]raclopride | D2R | 10 male | 10 male<br>2 female | ASD: 25(4)<br>Controls: 26(4) | DSM-V, ADOS-2 |
| *Noppari et al, 2024* | [11C]raclopride | D2R | 16 male | 24 male | ASD: 30(5)<br>Controls: 29(9) | DSM-V, ADOS-2 |
